# Ship-to-Shore Training for Active Deep-Sea Capacity Development

**DOI:** 10.1101/2023.03.11.531674

**Authors:** Kelsey Archer Barnhill, Beatriz Vinha, Alycia J. Smith, Daniëlle S.W. de Jonge, Daniela Y. Gaurisas, Roger Mocholí Segura, Pedro Madureira, Mónica Albuquerque, Veerle A.I. Huvenne, Covadonga Orejas, Vikki Gunn

## Abstract

Sailing on scientific expeditions as an early career researcher (ECR) offers the beneficial opportunity to gain field experience and training. However, the number of available berths to achieve the scientific goals of an expedition limits the number of onboard participants. Telepresence and remote learning can be utilised to increase the number of active participants, broadening the reach of capacity development. The 2021 iMirabilis2 expedition on board the Spanish Research Vessel *Sarmiento de Gamboa* used telepresence to virtually involve ECRs from several countries in deep-sea science. One year post-expedition, a survey of onshore participants was conducted to assess and quantify the effectiveness of the peer-to-peer ECR ship-to-shore scheme. During the expedition, live, interactive training via WhatsApp and Zoom was utilised by onshore ECRs more than traditional static, uni-directional methods of blog posts and pre-recorded videos. All respondents either agreed or strongly agreed that the scheme provided an inclusive and accessible platform to share deep-sea science. These results suggest similar schemes could be used to supplement shorter duration at-sea-training, used prior to a seagoing experience to better prepare ECRs, or to allow members of the science community unable to join an expedition in person to actively participate remotely, increasing inclusivity.

## Introduction

Scientific research expeditions are a key part of deep-sea science. When a project focused on the deep sea is funded, ship time is essential to carry out research on site. Cruises, particularly multi-day expeditions to the open ocean, are expensive (Ruth, 2006) and in many cases competition for ship time is strong, so most expeditions are necessarily multi-institutional, international and often multidisciplinary. In the short time onboard, data collection is optimized to occupy the entire time at sea, including during transit time to study sites when acquisition of bathymetry and measurement of physio-chemical parameters in the water column or atmosphere are commonplace. The data collected on these expeditions underpin large portions of project outputs and form the basis of multiple scientific publications.

Sailing on an expedition as an early career researcher (ECR) offers beneficial experience and training for career development through skill sharing and direct practice (Levin et al., 2019; Woodall et al., 2021). Expeditions not only offer hands-on experience on the practical aspects of data/sample collection but also reinforce the collaborative and multidisciplinary nature of marine science, offering ECRs the opportunity to work alongside scientists from different institutions, countries, and cultures. When participants experience a safe and inclusive working space during the expedition important ties and collaborations can be established through working and living together onboard whilst exchanging knowledge, ideas, and personal experiences (Amon, Filander, et al., 2022). Seagoing experience offers ECRs experience in team building, aptitude in handling onboard equipment, working in extreme conditions, and integrating skills in a new environment, which creates the need to develop adaptive management, creative problem-solving, and flexibility. This experience is often viewed as an essential part of the foundation for a career in at-sea marine science (Gunn & Thomsen, 2015). Ship size restrictions and expedition cost can cause restricted accessibility and participation for deep-sea science students and ECRs (Gerringer et al., 2023). Despite the number of berths onboard at times restricting the number of ECRs who can join, it is important to provide the next generation of marine scientists with appropriate field skills (Barnhill, 2022).

The UN Decade of Ocean Science for Sustainable Development recognises that to achieve their decade goals, ‘skills, knowledge, and technology for all’ will be required (IOC-UNESCO, 2021). Addressing existing global capacity and capability gaps in ocean science can only occur with advances in training methods and increased knowledge exchange across nations (Pendleton et al., 2020). Existing programs offering onboard ECR training such as NOAA’s Explorers in Training (https://oceanexplorer.noaa.gov/okeanos/training.html) and the Ocean Exploration Trust’s Science and Engineering Internships (https://nautiluslive.org/join/internship-program) utilise telepresence to further share their training to a wider at-home audience (Wishnak et al., 2022). First used in at-sea science by Dr. Robert Ballard in the JASON Project, telepresence allows meaningful scientific and educational ship-to-shore communications (Raineault et al., 2018). Telepresence can increase the number of remote participants in an ocean science expedition, broadening participation and transforming the field (Brennan et al., 2018; Cantwell et al., 2016; Marlow et al., 2017; Martinez et al., 2020). It also allows for direct and two-way training and communication that is lacking in the more traditional uni-directional methods of sharing offshore science, such as blog posts. The discovery aspect of deep-sea research is currently enjoying a surge of popularity with the general public and telepresence-enabled expeditions also offer an ideal outreach opportunity (Barnhill, 2022).

### iMirabilis2 Expedition Preparation

The EU Horizon2020 project iAtlantic (https://www.iatlantic.eu/) has science outreach and capacity-building of ECRs as one of its key objectives (Roberts et al., 2023). The iMirabilis expedition was originally planned as a multidisciplinary scientific expedition involving scientists from several countries bordering the Atlantic Ocean, with a strong training and outreach component and berths set aside for ECR partners based in Brazil and South Africa. However, challenges brought about by the Covid-19 pandemic necessitated significant re-planning and adjustments, resulting in a scaled-back expedition with a reduced number of participants (an experience that was shared by cruise organisers across the globe (Gallaudet et al., 2020)). Despite this, capacity development and outreach remained a mission priority, and additional effort and resources were directed to using virtual means to allow all those involved and interested in the expedition the ability to participate without physically boarding the ship.

In the lead up to the expedition, the first point of action was to recruit a dedicated onboard outreach liaison to join the ship, partnered with an onshore outreach liaison to create a core outreach team of two. Next, the team created an outreach and communication plan (found in (Orejas et al., 2022)), identifying the target audience, engagement opportunities beyond the iAtlantic network, and types of content to share during the expedition, following the considerations presented in Cooke et al. (2017). South African-based scientists within the One Ocean Hub project communicated their desire to the iAtlantic team to have trainings on deep-sea equipment created to help develop in-country capacity (Sink et al., 2021).

The onshore outreach liaison created a dedicated website (https://www.iatlantic.eu/imirabilis2-expedition/) for the iMirabilis2 expedition in the months leading up to the cruise (Figure 1). Prior to setting sail, pages were populated with background material and information to help set the context for the research being carried out on board, drawing on help from experts within the iAtlantic project team. This resource material included ‘Mission overview’, ‘Meet the team’, ‘Science on board’, ‘Equipment & technology,’ ‘Expedition blog’ and ‘Training & capacity building.’

**Figure 1.**
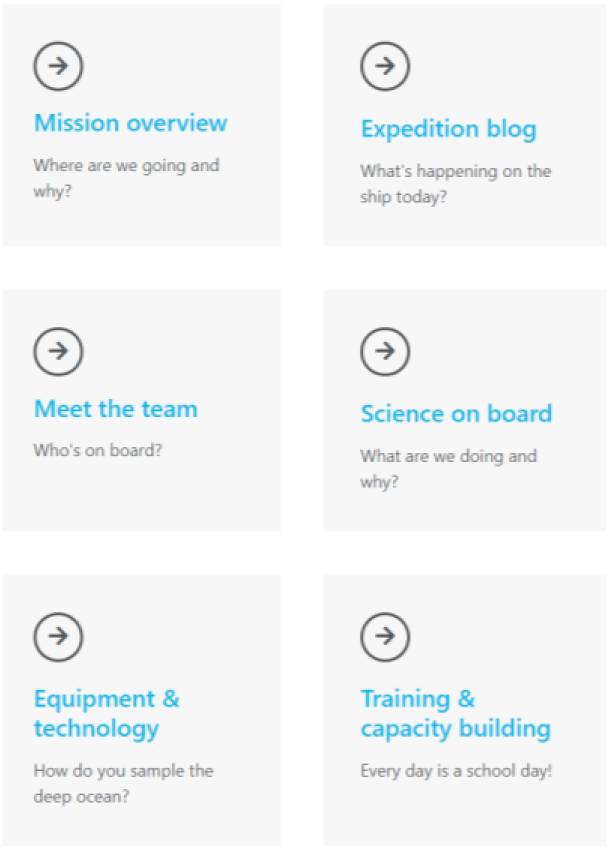
Screenshot of iMirabilis2 website’s home page (https://www.iatlantic.eu/imirabilis2-expedition/).

Interest in the outreach and capacity development components of the expedition was stimulated via presentations to the iAtlantic project team at a dedicated webinar, as well as to the wider deep-sea community through mailing lists such as Deep Ocean Stewardship Initiative (DOSI) (https://www.dosi-project.org/) and advertised on social media accounts. An online session titled ‘iMirabilis2: Deep Sea to Desktop’ was convened as an official satellite activity for the UN Ocean Decade Laboratory on ‘An Inspiring and Engaging Ocean’ to raise awareness in the wider scientific community and public about the forthcoming expedition coverage (https://www.iatlantic.eu/imirabilis2-expedition/deep-sea-to-desktop/). The expedition set sail with a commitment to shipboard and virtual capacity development and outreach, under the banner of ‘Deep sea to Desktop.’

### iMirabilis2 Expedition and Outreach

In July and August 2021, the iAtlantic iMirabilis2 expedition sailed onboard the Spanish Research Vessel *Sarmiento de Gamboa* (UTM-CSIC), heading to the Azores-Biscay Rise and Cabo Verde archipelago (Orejas et al., 2022). The expedition had several pieces of large equipment on board, namely the Task Group for the Extension of the Continental Shelf of Portugal’s (EMEPC) remotely operated vehicle (ROV) *Luso*, Heriot-Watt University’s Benthic Respirometer, Baited Camera, and Baited Trap Landers, and the National Oceanography Centre’s (NOC) Autonomous Underwater Vehicle (AUV) Autosub6000 equipped with the eDNA Robotic Cartridge Sampling Instrument, RoCSI. The expedition consisted of two legs, with Leg 0 led by EMEPC and Leg 1 led by the Spanish Institute of Oceanography (IEO), a partner from the iAtlantic project (Figure 2). Leg 0 focused on acquiring seafloor bathymetry and geological and biological samples from the Northern Azores-Biscay Rise to characterise the environment and better understand North Atlantic Basin evolution. Leg 1 focused on exploring benthic ecosystems in the abyssal plains off Brava Island and the continental shelves, slopes and bathyal zone off Brava and Fogo Islands, with specific interest in the islands’ slopes and nearby Cadamosto Seamount. Training and outreach activities were an important component of both expedition legs.

**Figure 2.**
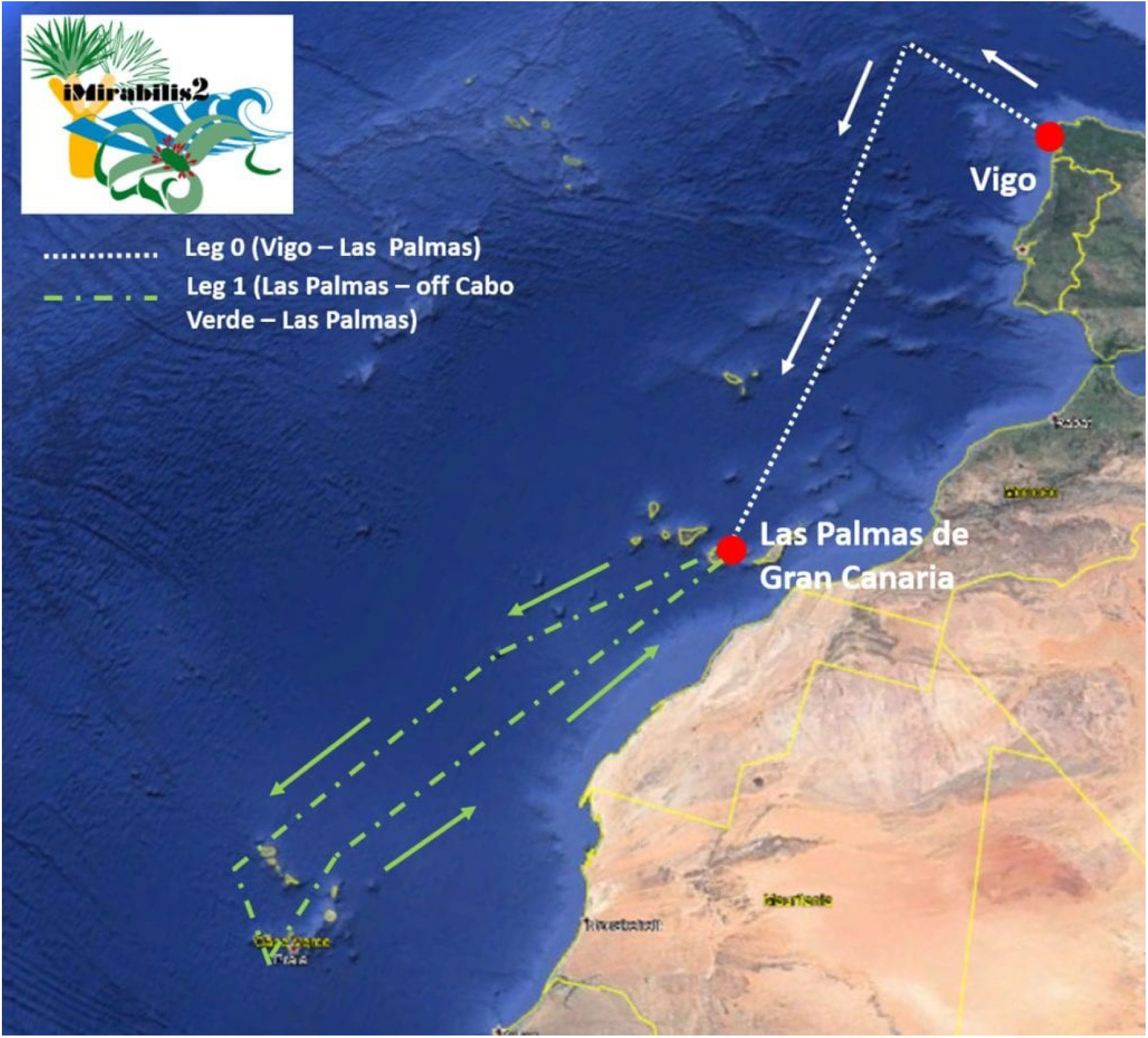
iMirabilis2 Expedition Route

During the expedition, the onshore outreach liaison was responsible for onshore support while the onboard outreach liaison led offshore content creation. Content created onboard the ship was sent to shore to be edited, formatted, and uploaded to the website. The onboard outreach liaison received support from the whole expedition party, specifically from other ECRs in producing training and outreach material. The two core team members used email and WhatsApp to communicate with each other during the expedition. Files too large for email transmission, such as multiple high resolution photographs and videos, were sent via the MyAirBridge platform (https://www.myairbridge.com/).

Internet from an onboard very small aperture terminal (VSAT) was utilised to provide live outreach and training from the expedition. When fully functioning, the onboard bandwidth ranged from 4 to 7 MB/s. Despite significantly slower internet speeds than the average European household (the EU Digital Agenda to Europe aimed for universal coverage of 30 MB/s by 2020) (European Comission, 2014), internet speeds were sufficient for Zoom calls, which require just 600 KB/s (Chang et al., 2021). To support internet stability during Zoom video calls, all ship bandwidth was diverted to a single computer (aside from access at the bridge for safety reasons). Unfortunately, bandwidth was not sufficient to livestream ROV dives, which require faster internet speeds, such as NOAA’s *Okeanos Explorer*’s rates of 40 MB/s (Gallaudet et al., 2020). Additional internet privileges were also given to the onboard outreach liaison to send larger files to shore. These internet privileges were made possible due to the high priority placed in capacity development during the cruise planning stage.

Existing iAtlantic social media accounts were used for outreach purposes, which has been noted to inspire researchers in developing countries and indirectly supports capacity development (Sink et al., 2021). Updates from the ship including images, videos, and blog posts were shared via Twitter (@iAtlanticEU), YouTube (iAtlantic H2020), Instagram (@iatlanticeu), and Facebook (https://www.facebook.com/iAtlanticEU). One Reddit AMA (ask me anything) event was scheduled to expand reach beyond iAtlantic social media followers (https://www.reddit.com/r/askscience/comments/pcl2nr/askscience_ama_series_were_marine_scientists/). The majority of social media updates from iAtlantic accounts were made directly from the ship. Partners such as EMEPC and NOC used their websites and social media accounts to raise the expedition profile and share cruise information.

During expedition coverage the @iAtlanticEU Twitter account gained 150 new Twitter followers, experienced a 150% impression increase, and had a 38% increase in profile visits. As the iAtlanticEU Facebook page was previously not updated as frequently as the Twitter account, the regular coverage during the expedition led to an increase in reach of over 2000%. Social medial metrics and impressions were further increased due to partners sharing original content related to the expedition as well as content produced from iAtlantic Project accounts. The iAtlantic website experienced an uptick in the number of daily views in the time leading up to and during the iMirabilis2 Expedition. Increased engagement on the iAtlantic social media accounts during the expedition, shown by the increase in followers and engagement across apps, suggests there is a strong interest from the public in following deep-sea expeditions.

During the iMirabilis2 expedition 36 blog posts were published, ∼100 original Tweets were shared, 17 videos were uploaded to the YouTube Channel, and 23 Facebook posts and 17 Instagram posts were created. The onboard outreach liaison authored 22 blog posts, the onshore outreach liaison wrote four of the posts, and the remaining 10 were guest-authored by members of the onboard science team. Blog post topics ranged from scientific to life onboard.

Scientific topics included:

- ROV dive recaps focused on images and descriptions of the studied habitats
- How we process and store samples from the ROV
- The annotation process during a dive
- The seafloor mapping process
- How seabird surveys are conducted
- How the new RoCSI collects eDNA samples
- How the benthic respirometer lander conducts seafloor experiments
- Images from the baited camera trap deployments
- Descriptions of the catch from the baited trap
- Initial results from the onboard sediment incubation experiment

Posts written on life onboard included:

- Items we pack for an expedition
- Last-minute preparations completed the day prior to boarding
- Food we eat onboard
- First impressions of the ship
- Safety drill experience
- A science team member’s first at-sea experience

Training videos created and shared during the expedition included:

- What is the role of a science team member during an ROV dive?
- How to process ROV samples
- How to prepare for AUV missions
- An introduction to the equipment (Benthic Landers, AUV, ROV, CTD)
- A description of incubation experiments

Additional videos created and shared included:

- A camera lander time-lapse
- ROV deployments
- ROV footage highlights
- A guided tour of the research vessel

All videos were created using an iPhoneSE, Rode Smartlav+ Lavalier microphone for smartphones with a wind shield, and a smartphone tripod. Videos were edited using the open-source software Shotcut (https://shotcut.org/).

### Onboard Capacity Development

The flagship capacity development effort on the expedition was the ‘Ship to Shore Buddies’ scheme, where ECRs onboard shared their experiences in a personalised way to their peers onshore. Efficient, constant, and quick real-time interactions between the onboard and onshore ECRs occurred via a dedicated WhatsApp group which had over 250 messages by the end of the expedition. A one-hour Zoom call between the ECRs was scheduled each week to share expedition progress and answer any questions about a researcher’s normal day at sea and the work carried out onboard. In total, the onboard and onshore ECRs met up for five Zoom interactions. Onshore ECRs had the opportunity to ask questions about any aspect related to the expedition. Topics discussed during Zoom interactions included life onboard, dealing with seasickness and homesickness, internet and phone connection on board, and the living quarters. The main focus of calls was to explain how the team work with the various types of equipment as well as what data was collected before discussing preliminary findings in depth, collaborating together to understand the data. Training videos on the most commonly used equipment and methods were also shared on these calls before they were made available to the general public via YouTube. Following the calls, all training materials were published and remain publicly available on the expedition website, as well as on the iAtlanticEU YouTube account. Creating deep-sea education materials like the training videos help share scientific advances and best practices for at-sea equipment use to students and ECRs (Levin et al., 2019). Eighteen marine science ECRs participated as onshore ECRs, joining the scheme from Brazil, Cabo Verde, Ghana, Portugal, and South Africa. Scheme insights from some onshore ECRs were shared on the expedition blog at the end of the cruise (https://www.iatlantic.eu/expedition_blog/ship-to-shore-a-virtual-expedition/).

One year post-expedition, a survey for the onshore participants was created to understand and quantify the effectiveness of the ship-to-shore buddies scheme. The survey was created in Jisc and consisted of 17 close-ended questions and five open-ended questions. Close-ended questions were phrased as: ‘To what extent do you agree or disagree with the following statements?’ With the response options being: ‘strongly agree’, ‘agree’, ‘neither agree nor disagree’, ‘disagree’, and ‘strongly disagree.’ All close-ended questions were mandatory for respondents to complete prior to survey submission while all open-ended questions were optional. The survey was sent out via email to all 18 shore-based ECRs. It remained open for three weeks with one reminder email sent in the final week of the survey remaining open to new responses. The response rate was 55% with 10 ECRs completing the survey.

### Impact of the Ship-to-Shore Buddy Scheme

Survey results were used to assess the effectiveness of the ship-to-shore buddies scheme from the point of view of the recipients, which is important to consider for equitable ocean science capacity development (Harden-Davies, Amon, Vierros, et al., 2022). Close-ended questions were used to look at how the onshore ECRs engaged with, were impacted by, and experienced the scheme (Table 1). The majority of the respondents reported watching the videos (60%), reading the blog (70%), joining Zoom calls (80%), and following the WhatsApp chat (100%). The majority also found the Zoom calls and WhatsApp groups helpful (80 and 90% respectively). Sixty percent of respondents reported learning something useful to developing their research and referring to materials after the end of the expedition. Half of the participants reported using the knowledge and/or contacts gained during the scheme in the past year. Regarding the experience of participating in the scheme, all respondents reported either agreeing or strongly agreeing (70% and 30% respectively) that the scheme allowed an inclusive and accessible way to share deep-sea science.

**Table 1.** Close-ended survey results. Respondents were asked ‘To what extent do you agree or disagree with the following statements?’

|  | Strongly Agree | Agree | Neither Agree nor Disagree | Disagree | Strongly Disagree |
| --- | --- | --- | --- | --- | --- |
| I felt actively engaged in the iMirabilis2 cruise | 3 | 5 | 2 | - | - |
| I learned about conducting at-sea research in a personalised way | 3 | 7 | - | - | - |
| In the past year, I have used the knowledge and/or contacts I gained during the scheme | 1 | 4 | 3 | 2 | - |
| I acquired knowledge useful to developing my research/thesis/study during iMirabilis2 | 3 | 3 | 4 | - | - |
| The scheme was accessible | 6 | 4 | - | - | - |
| I felt like a part of an early career researcher network | 4 | 6 | - | - | - |
| I would be interested in virtually following along with another future expedition | 6 | 4 | - | - | - |
| I would recommend this scheme to fellow early career researchers | 8 | 1 | 1 | - | - |
| I read the iMirabilis2 website blog posts | 3 | 4 | 1 | 1 | 1 |
| I watched the video training material either on the iMirabilis2 website or YouTube | 5 | 1 | 2 | 2 | - |
| I referred to the online materials after the cruise ended | 1 | 5 | 2 | 1 | 1 |
| I joined in on Zoom calls during the cruise | 7 | 1 | 1 | 1 | - |
| I found the Zoom calls helpful | 7 | 1 | 2 | - | - |
| I followed the WhatsApp group chat during the cruise | 8 | 2 | - | - | - |
| I found the WhatsApp group helpful | 7 | 2 | 1 | - | - |
| The buddy scheme could be used to supplement at-sea training to prepare early career researchers for a cruise | 7 | 3 | - | - | - |
| This scheme could be used to promote more inclusive at-sea experiences | 7 | 3 | - | - | - |

Major themes were identified across the respondents’ answers in the open-ended questions using NVivo software. Themes identified were: learning about deep-sea science, methods and equipment, feeling part of an ECR network, using the scheme to prepare for future research, learning about life onboard, and how participants could replicate a similar scheme on future expeditions. To share and demonstrate the experience the onshore ECRs reported in their survey responses, one quote from each theme is presented below.

#### Deep-Sea Science and Equipment

‘The experience has taught me about sampling equipment that we do not currently have access to in South Africa. Over the past year I have used this knowledge to assist with the writing of project and equipment proposals.’

#### ECR Network

‘Getting to know other early-career researchers and receiving their impressions of what was (sic) like to be on an expedition first-hand was definitively the most useful aspect of the buddy scheme.’

#### Prepare for Future Research

‘The experience helped me understand the protocols and workflows of research (sic) cruise outfitted with state of the art underwater camera platforms. I have incorporated aspects of the of the (sic) workflows into my own when at sea. It has also helped me earmark research areas of interest that I was unaware of an (sic) would like to explore.’

#### Life Onboard

‘It was surely an enriching experience, where I got to know some of the details of what a research expedition is like and had the opportunity to learn directly from other researchers.’

#### Replicate a Similar Scheme

‘The platform was innovative and engaging. It has prompted me to establish something similar whenever I go to sea as a means to engage fellow scientists onshore.’

## Discussion

Establishing outreach and capacity development as priorities from the outset of expedition planning allows it to develop in tandem with scientific goals and improves the chances of a successful scheme. The iMirabilis2 ECR ship-to-shore buddy program is by no means a direct replacement for seagoing experience, however our results suggest it could be used to supplement shorter duration at-sea training or used prior to a seagoing experience to better prepare ECRs. While onshore participants who completed the survey provided largely positive feedback, this one-off experience alone is not sufficient to ensure lasting capacity, which can only be achieved through long-term partnerships (Amon, Rotjan, et al., 2022; Harden-Davies, Amon, Chung, et al., 2022). Restrictions caused by Covid-19, which initially threatened to decrease opportunities for ECR training and capacity development, especially for participants from the countries most affected during the pandemic, ultimately broadened the reach through virtual learning. Within the UN Decade of Ocean Sciences, there is a push to make deep-sea science more inclusive (Howell et al., 2020), and projects led by European countries with high levels of deep-sea capacity can help facilitate global access to the deep-sea (Bell et al., 2022). While Sink et al. (2021) identify lack of technology and vessel access as major challenges towards developing deep-sea research capacity in South Africa, they also identify training and exposure as needs. This virtual training method could be implemented again in the future, to broaden access and improve accessibility for science community members unable to join expeditions. This result is supported by one ECR onshore respondent who shared:

‘My recommendation will be that these scheme (sic) should be given more publicity so that it is a default part of every scientific cruise. I believe it will go a long way in providing the needed experience for ECRs especially in developing countries where such opportunities are scarce.’

Scientific expeditions often share their experiences and findings to a wider audience via one-way communications such as a blog (Bingham et al., 2015) or through social media posts (which are often shared after an experience has taken place), but the majority of engagement during the iMirabilis2 expedition came from live interactions. Securing the bandwidth required to operate a telepresence-enabled cruise comes at significant expense, but allows for a more just and inclusive deep-sea expedition (Cantwell et al., 2020). Scientific capacity development and peer-to-peer learning on the iMirabilis2 cruise occurred in real time over the WhatsApp group chat and Zoom calls. Both WhatsApp and Zoom are commonly-used free or low-cost tools which are easily accessible and do not require additional training to utilise. Both technologies can be easily implemented on other expeditions, including those with smaller budgets and less bandwidth. For expeditions with sufficient bandwidth for livestreaming ROV dives and other onboard scientific operations, incorporating onshore ECR-led research into the cruise plan is another successful way to develop capacity (Pallant et al., 2018; Stephens et al., 2016).

The WhatsApp group likely also broadened the scheme’s reach, as participation was not limited by time zones and schedules. The onboard ECRs were accessible to onshore participants around the clock due to 24-hour shift patterns, including when bandwidth was unreliable as WhatsApp messages can still be sent and received on slower networks. This more informal method of communication also allowed the international ECRs to form networks, collaborations and friendships. To increase accessibility for onshore ECRs to participate, future projects could consider offering technical or financial assistance to ensure internet connections allow individuals to participate remotely if required.

Creating an effective outreach and training program is a large task that should not be placed on someone whose main role onboard is scientific or technical. This was recognised as early as 2004 with Sautter (2004) recommending cruise leaders write a marine science educator into grants to share at-sea science with the public. The International Ocean Discovery Program (IODP) Onboard Outreach Officer program recognises this need and hires a paid science communicator to join each expedition to create this content (Garnsworthy & Kurtz, 2019). Having an onshore outreach liaison provide support to the onboard outreach liaison allowed for sharing of workloads. To work around reduced internet speeds during the expedition, the onboard team member was solely responsible for sending created content to the onshore outreach liaison, who was responsible for formatting and sharing the content via the website. Balancing time on an expedition is often difficult for cruise participants. When incorporating an outreach member in an expedition, ensuring a clear prioritisation order for science vs outreach tasks onboard is crucial for time management. In this case, the onboard outreach liaison was also a member of the science team but was made aware that outreach was the main focus of their role. The interactive WhatsApp and Zoom calls also took less time and effort for the onboard participants during the expedition than creating blog posts and videos, an important factor to consider when onboard participants have other commitments. Being included in the scientific team was advantageous for the outreach liaison, who was able to speak with experience on the science conducted onboard, as they played an active role in several tasks. Having an outreach liaison who came from a scientific background was also beneficial when describing research they were not involved in, as they were familiar with many concepts and the overall field of deep-sea science. In addition to being able to share scientific concepts accurately, it is important for the onboard outreach liaison to be able to translate offshore science effectively to the desired audience, using language free of scientific jargon (Westnedge & Dallimore, 2014).

The outreach team identified having time dedicated to completing outputs after the expedition as a key way to improve future similar schemes. Following the expedition end and momentum gained during the cruise coverage, the outreach team and ECRs onboard had unedited videos which were never completed and shared, as well as ideas for more live and interactive virtual trainings on ROV annotations. Showing students and ECRs recorded ROV dives and allowing them to formulate individual research questions surrounding the video is another method which can expand deep-sea accessibility (Gerringer et al., 2023). Ensuring dedicated time to completing outputs post-expedition would ensure these projects were finished and implemented, further improving two-way capacity development opportunities for onshore participants.

The more traditional forms of shipboard outreach, such as blog posts and educational material, were not utilised as much by the remote ECRs as compared to the newer and more interactive methods. All survey respondents reported following the WhatsApp group with eight out of 10 participants reporting also joining in on a Zoom call, while just six of the 10 respondents reported watching video training materials and seven out of the 10 reported reading blog posts. One survey respondent who only participated in the WhatsApp chat still reported feeling actively engaged in the expedition. Real-time ship-to-shore interactions have previously been identified as key to student learning processes, as they allow students to connect to the wider deep-sea community (Sánchez et al., 2020). We suggest these real-time personalised interactions drove the positive outcome of the scheme.

## Supporting information

Supplemental Data: Survey

## Acknowledgements

The authors would like to thank Professor J. Murray Roberts, Christine Gaebel, Hannah Elise Barker, the entire team at Unidad Technología Marina (UTM, CSIC), the team at Estrutura de Missão para a Extensão da Plataforma Continental (EMEPC), as well as the entire technical and scientific team and crew onboard the Spanish R/V *Sarmiento de Gamboa* (UTM-CSIC) for the iMirabilis2 Expedition. Thank you to the Spanish Ministry of Science and Innovation for providing ship time. Kelsey Archer Barnhill thanks the Deep-Sea Biology Society for funding her cruise participation. This project has received funding from the European Union’s Horizon 2020 research and innovation programme under grant agreement No 818123 (iAtlantic). This output reflects only the authors’ views and the European Union cannot be held responsible for any use that may be made of the information contained therein.

## Data Availability

Data supporting this study cannot be made available due to privacy concerns for survey participants. Respondents were told access to survey results would be limited to the researchers authoring this paper.

## Ethics Statement

The work completed in this article was reviewed and approved by the University of Edinburgh School of GeoSciences Research Ethics & Integrity Committee (reference number 2022-636).

