## Supplemental Data: Survey for "Ship-to-Shore Training for Active Deep-Sea Capacity Development"

### iMirabilis2 Ship-to-Shore Buddy Scheme Impacts

#### Information and Consent

**Information Sheet**

*Research project title*: iMirabilis2 Ship-to-Shore Buddy Scheme Impacts

*Research investigator*: Kelsey Archer Barnhill

Kelsey led the on board ECR Buddy Scheme and helped organize it alongside Vikki Gunn. Contact details of research investigator:

*Other Researchers may be involved in this project. Their names and roles are:*

Vikki Gunn, Director, Seascape Consultants Ltd.

Beatriz Vinha, PhD Student, University of Salento

Danielle de Jonge, PhD Student, Heriot-Watt University

Alycia Smith, PhD Student, Heriot-Watt University

Daniela Yepes Gaurisas, PhD Student Universidade Federal do Espírito Santo

*About the Project*

In July 2021, the iAtlantic iMirabilis2 cruise set sail with a commitment to shipboard and virtual capacity building and outreach under the catchphrase 'Deep-sea to Desktop.' The flagship effort of capacity building on the cruise was the 'Ship to Shore Buddies' scheme where ECRs onboard shared their experiences in a personalised way to their peers onshore in real time.

The aim of this survey is to quantify the success of this peer-to-peer learning and networking program. This survey will better help guide future on board outreach opportunities in deep-sea science, broadening participation and creating strong ECR networks in the field.

*What are the risks involved in this study?*

We don’t anticipate that there are any risks associated with your participation, but you have the right to stop responding to the survey or withdraw from the research at any time.

• Access to the survey results will be limited to Kelsey Archer Barnhill and academic colleagues and researchers with whom he might collaborate as part of the research process

• Any survey content, or direct quotations from the survey, that are made available through academic publication or other academic outlets will be anonymized so that you cannot be identified, and care will be taken to ensure that other information in the survey that could identify yourself is not revealed

• Any variation of the conditions above will only occur with your further explicit approval

*How will my data be used?*

All or part of the content of your survey may be used:

· In academic papers, policy papers or news articles

· On our website and in other media that we may produce such as spoken presentations

· On other feedback events

*What are your rights as a participant?*

Taking part in the study is voluntary. You may choose not to take part or subsequently cease participation at any time.

*Will I receive any payment or monetary benefits?*

You will receive no payment for your participation. The data will not be used by any member of the project team for commercial purposes. Therefore you should not expect any royalties or payments from the research project in the future.

*For more information*

This research has been reviewed and approved by the Edinburgh University Research Ethics Board. If you have any further questions or concerns about this study, please contact:

Kelsey Archer Barnhill

*What if I have concerns about this research?*

If you are worried about this research, or if you are concerned about how it is being conducted, you can contact the Chair of the GeoScience Ethics Committee, University of Edinburgh, Drummond St, Edinburgh, EH8 9XP (or email at ).

**Survey Consent**

Thank you for agreeing to answer questions as part of the above research project. Ethical procedures for academic research undertaken from UK institutions require that participants explicitly agree to participate and agree to how the information contained in their responses will be used. Your consent is necessary for us to ensure that you understand the purpose of your involvement and that you agree to the conditions of your participation. Would you therefore read the above information sheet and then certify that you approve the following:

Your answers will be analysed by the research investigator and any summary of your responses, or direct quotations that are made available through academic publication or other academic outlets will be anonymised.

Prior to your response, you agree that:

1. You are voluntarily taking part in this project and understand that you are not required to take part;

2. Any extracts from your responses may be used as described above in this research project;

3. You have read the Information section;

4. You don't expect to receive any benefit or payment for your participation;

5. You understand that you are free to contact the researcher with any questions you may have in the future.

**Informed Consent** I confirm that I have read and understood the Participant Information Sheet for the above study. I understand that my anonymised data will be stored for a minimum of 5 years and may be used in future ethically approved research. 1. By selecting 'Yes'  I consent to participate in the survey ✱ *Required*

- ◯ Yes
- ◯ No

### Questions 1/10

To what extent do you agree or disagree with the following statements? ✱ *Required*

Please don't select more than 1 answer(s) per row.

Please select at least 17 answer(s).

| | Strongly agree | Agree | Neither agree nor disagree | Disagree | Strongly disagree |
|---|---|---|---|---|---|
| I felt actively engaged in the iMirabilis2 cruise. | ☐ | ☐ | ☐ | ☐ | ☐ |
| I learned about conducting at-sea research in a personalised way. | ☐ | ☐ | ☐ | ☐ | ☐ |
| In the past year, I have used the knowledge and/or contacts I gained during the scheme. | ☐ | ☐ | ☐ | ☐ | ☐ |
| I acquired knowledge useful to developing my research/thesis/study during iMirabilis . | ☐ | ☐ | ☐ | ☐ | ☐ |
| The scheme was accessible. | ☐ | ☐ | ☐ | ☐ | ☐ |
| I felt like a part of an early career researcher network. | ☐ | ☐ | ☐ | ☐ | ☐ |
| I would be interested in virtually following along with another future expedition. | ☐ | ☐ | ☐ | ☐ | ☐ |

| | | | | | |
|---|---|---|---|---|---|
| I would recommend this scheme to fellow early career researchers. | ☐ | ☐ | ☐ | ☐ | ☐ |
| I read the iMirabilis2 website blog posts. | ☐ | ☐ | ☐ | ☐ | ☐ |
| I watched the video training material either on the iMirabilis2 website or Youtube. | ☐ | ☐ | ☐ | ☐ | ☐ |
| I referred to the online materials after the cruise ended. | ☐ | ☐ | ☐ | ☐ | ☐ |
| I joined in on Zoom calls during the cruise. | ☐ | ☐ | ☐ | ☐ | ☐ |
| I found the Zoom calls helpful. | ☐ | ☐ | ☐ | ☐ | ☐ |
| I followed the WhatsApp group chat during the cruise. | ☐ | ☐ | ☐ | ☐ | ☐ |
| I found the WhatsApp group helpful. | ☐ | ☐ | ☐ | ☐ | ☐ |
| The buddy scheme could be used to supplement at-sea training to prepare early career researchers for a cruise. | ☐ | ☐ | ☐ | ☐ | ☐ |

| | | | | | |
|---|---|---|---|---|---|
| This scheme could be used to promote more inclusive at-sea experiences. | ☐ | ☐ | ☐ | ☐ | ☐ |

In the past year, how has the experience impacted or helped your work/career?

In your opinion, what was the most useful aspect of the buddies scheme?

In a few sentences, how would you explain your experience participating in the buddies scheme?

What suggestions would you make to improve the ship-to-shore buddies scheme?

Do you have any additional comments?

### Final page

Thank you for taking the time to answer this survey. Your responses have been recorded. Any outputs associated with the results of this survey will be shared with you.
